# Beta-subunit-eliminated eHAP expression (BeHAPe) cells reveal new properties of the cardiac voltage-gated sodium channel

**DOI:** 10.1101/2023.07.08.548226

**Authors:** Annabel Y Minard, Colin J Clark, Christopher A Ahern, Robert C Piper

## Abstract

Voltage-gated sodium (Na_V_) channels drive the upstroke of the action potential and are comprised of a pore-forming α-subunit and regulatory β-subunits. The β-subunits modulate the gating, trafficking, and pharmacology of the α-subunit. These functions are routinely assessed by ectopic expression in heterologous cells. However, currently available expression systems may not capture the full range of these effects since they contain endogenous β-subunits. To better reveal β-subunit functions, we engineered a human cell line devoid of endogenous Na_V_ β-subunits and their immediate phylogenetic relatives. This new cell line, β-subunit-eliminated eHAP expression cells (BeHAPe), were derived from haploid eHAP cells by engineering inactivating mutations in SCN1B, SCN2B, SCN3B, SCN4B, MPZL1, MPZL2, MPZL3, MPZ and JAML. In diploid BeHAPe cells, the cardiac Na_V_ α-subunit, Na_V_1.5, was highly sensitive to β-subunit modulation and revealed that each β-subunit and even MPZ imparted unique gating properties. Furthermore, combining β1 and β2 with Na_V_1.5 generated a sodium channel with hybrid properties, distinct from the effects of the individual subunits. Thus, this approach revealed an expanded ability of β-subunits to regulate Na_V_1.5 activity and can be used to improve the characterization of other α/β Na_V_ complexes.

## Introduction

Voltage-gated sodium (Na_V_) channels drive the upstroke of the action potential in excitable cells, thereby generating the electric signals that underlie behavior, sensation, muscle contraction and mobility (1-4). Na_V_ channels are composed of one pore-forming α-subunit and one or more regulatory β-subunits. The α-subunits are large (∼250 kDa) proteins with 24 transmembrane segments that are arranged in four domains (DI-DIV) to form a central sodium selective and conducting pore. The β-subunits are not required for conduction, but instead function to regulate the α-subunit. Their functions include trafficking and retention of the α-subunit at the plasma membrane, altering the pharmacology of the α-subunit, and altering the voltage-dependent gating properties of the α-subunit (2,5,6). These functions are crucial for proper electrical conduction, since even relatively small changes in the gating equilibria and rates between closed, open, and inactivated states can cause electrical dysfunction (7,8). There are ten α-subunit isoforms and five Na_V_ β-subunit isoforms. The Na_V_ β-subunits, β1-4, are single-pass membrane proteins, with an extracellular immunoglobulin (1g)-like domain. These β-subunits are encoded by the genes SCN1B, SCN2B, SCN3B, and SNC4B. A fifth Na_V_ β-subunit, B1b, generated from a splice variant of SCN1B, is expressed as a secreted protein containing the Ig-like domain. The predominant Na_V_ channel in the heart is Na_V_1.5, which forms complexes with β1-β4 (9-11). Loss of Na_V_1.5 in mice is lethal *in utero* while loss of β1-β3 causes cardiac arrhythmia (12-14). Furthermore, clinical variants in Na_V_1.5 or β1-β4 are associated with cardiac arrythmias and disorders such as Brugada Syndrome (BrS) and Long QT syndromes (LQT) (15,16). Despite the importance of Na_V_1.5 and β-subunits, precisely and systematically defining the effect of β-subunits and their diseases variants, on the electrophysiology of Na_V_1.5 has been challenging due in part to the limitations of currently available expression systems (17).

The electrophysiological properties of Na_V_ channels are routinely determined in heterologous systems, such as HEK293T, CHO and COS cells. These systems lack endogenous voltage-gated ion channels, allowing for the precise measurement of action-potentials generated by ectopically expressed α-subunits. However, they express endogenous β-subunits, which could alter α-subunit properties and interfere with the response to ectopically expressed β-subunits. To circumvent this issue *Xenopus laevis* oocytes are sometimes used since they have minimal endogenous β-subunit expression (18,19). But because *X. laevis* oocytes require a low incubation temperature (18° C), they are not ideal for the study of disease variants. In addition to endogenous β-subunits, the presence of phylogenetic relatives of β-subunits may also modify a channel-of-interest. The β-subunits belong to a sub-family of proteins (β/MPZ) that includes MPZ, MPZL1, MPZL2, MPZL3 and JAML (20,21). Although not implicated in directly regulating Na_V_ channels, these proteins may regulate the channel in cultured cells if they share biophysical properties with β-subunits. Indeed, the sequences of β1 and β3 are as similar to the MPZ subfamily as they are to β2 and β4 (20). Thus, an ideal expression system for studying Na_V_ β-subunit isoforms and their disease-associated alleles should lack an endogenous β/MPZ family.

In this study we used CRISPR-Cas9 gene-editing to generate **β**-subunit-**e**liminated e**HAP e**xpression (BeHAPe) cells, a cell line devoid of the β/MPZ gene family. This cell line allows β-subunits to be expressed as the sole β-subunit family member without interference from endogenous β-subunits. We used these cells to assess the effects of β-subunits on the gating properties on the predominant cardiac α-subunit Na_V_1.5. We found that in BeHAPe cells Na_V_1.5 was highly responsive to β-subunit effects and that co-expression with β-subunits generated a repertoire of Na_V_1.5 channels with unique gating properties. These findings demonstrate that a β/MPZ null cell model improves the characterization of the electrophysiological properties of α/β Na_V_ complexes.

## Results

### Generation of BeHAPe cells

To better reveal how β-subunits modulate the gating properties of α-subunits, we generated a cell line devoid of the β-subunit family (SCN1B, SCN2B, SCN3B, SCN4B) and their phylogenetic relatives (MPZ, MPZL1, MPZL2, MPZL3 and JAML). We deleted these genes from human haploid (eHAP) cells. eHAP cells are fibroblast like cells that do not produce endogenous voltage-gated sodium currents. These cells are also haploid, simplifying mutagenesis and genotyping. Previous RNAseq experiments (Accession: SRX655513, SRX65551) show that this cell line has low to moderate expression of β/MPZ family members (22). No expression of SCN2B or JAML was detected, whereas SCN1B, SCN3B, SCN4B, MPZ, MPZL1, MPZL2, and MPZL3 had FPKM values of 0.70, 0.20, 0.016, 2.15, 3.40, 0.01, and 0.30, respectively, comparable to those in HeLa and HEK293 cells (SRR1567907, SRR5011299).

We first integrated an FRT site into eHAP cells to allow for site-directed recombination in future studies (Figure 1A). The FRT site was encoded in the pQCXIP FRT GFP-Neo vector, which was transduced into eHAP cells by retrovirus. We mapped the integration site to chromosome 19, in between GPR108 and TRIP10 genes. The introduced locus expressed EGFP fused to neomycin resistance gene (eGFP-Neo^R^) under a CMV promoter (pCMV). An FRT site was placed between the start codon and eGFP. Later, in the second round of CRISPR-Cas9 deletions, we deleted eGFP-Neo^R^, leaving the CMV promoter and FRT site available for optional use.

**Figure 1.**
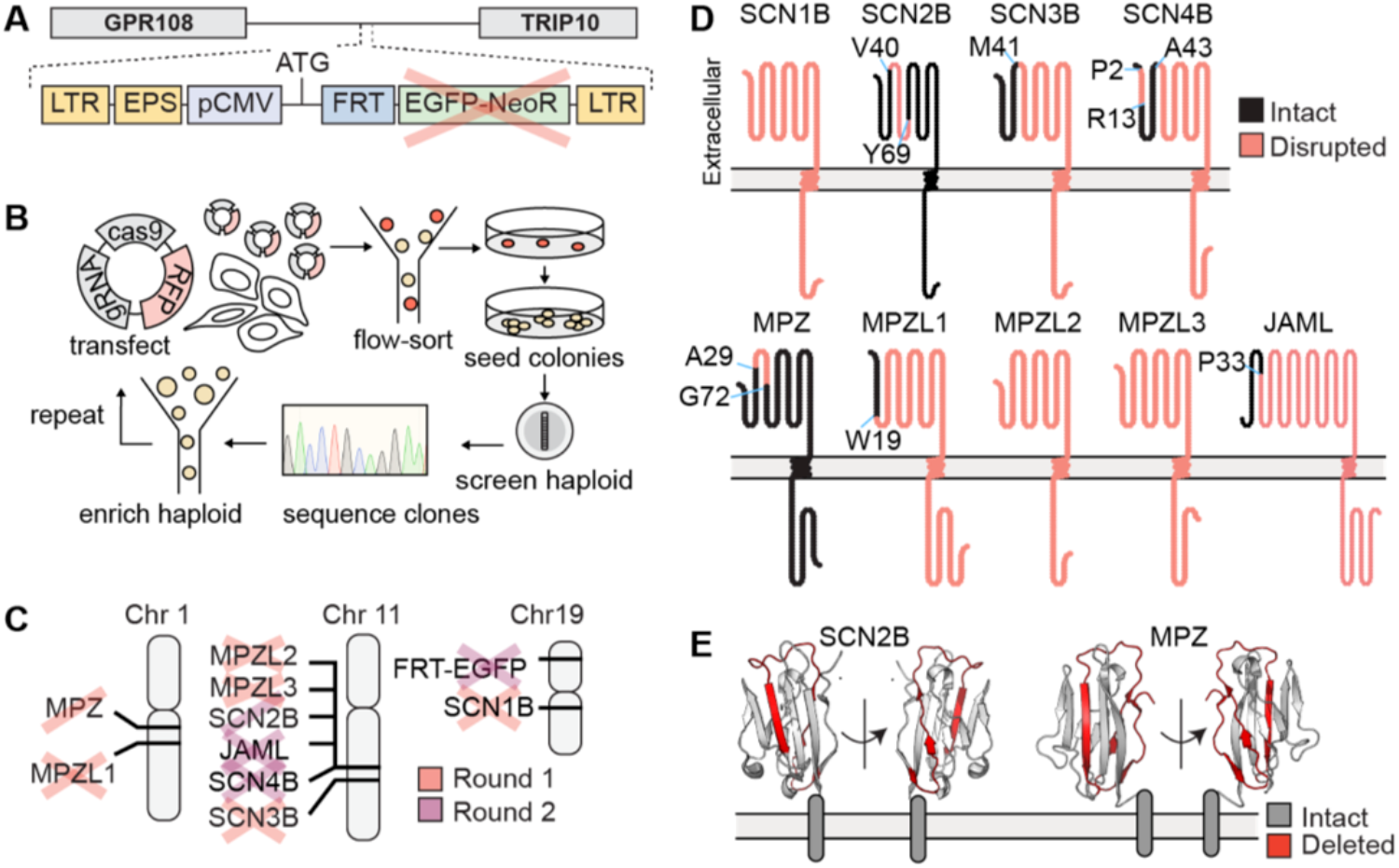
Generation of BeHAPe cells by two rounds of CRISPR-Cas9 deletion of β-subunits and their phylogenetic relatives. **(A)** eHAP-FRT cells have an FRT integration site between GPR108 and TRIP10. EGFP-NeoR was originally expressed from this locus but deleted in the second round of CRISPR-Cas9 deletions. **(B)** eHAP-FRT cells were engineered into BeHAPe cells with two rounds of CRISPR-Cas9. Multiple gRNA, Cas9 and fluorescent marker encoding plasmids were transfected into the eHAP-FRT cell-line. Transfected cells were enriched for by flow-sorting for fluorescent cells and serially diluted onto 10 cm dishes. Clones were screened to verify haploidy and then genotyped. **(C)** Genes marked with a cross were disrupted in BeHAPe cells, while a single-line indicates the gene was edited but not with a frame-shifting mutation. Whether a gene was targeted in round 1 or round 2 are indicated. **(D)** Amino acid residues that are no-longer encoded for in the edited genes are shown. Protein sequence was visualized with Protter (62). **(E)** Residues deleted in SCN2B and MPZ genes encoded for a β-strand in the Ig domain (SCN2B PDB:5FDY and MPZ PDB: 30AI).

We next disrupted the β/MPZ family in eHAP-FRT cells in two rounds of CRISPR-Cas9 gene editing (Figure 1B). For these experiments, Cas9 and gRNA encoding plasmids, co-expressing fluorescent reporter proteins, were transiently transfected into cells. To edit nine genes in two rounds of CRISPR-Cas9, we introduced multiple gRNA per gene and targeted multiple genes in each round. To improve the efficiency of gene edits we enriched for haploid cells prior to transfecting gene-editing reagents, which was necessary as eHAP cells diploidize. After transfection, we also enriched for fluorescent transfected cells by FACS because transfection efficiency was roughly 30%. We identified knockouts by PCR amplifying and Sanger sequencing the targeted genes. Because eHAP cells are haploid, they possess only one copy of the targeted genes. Knockouts were selected from cells with indel mutations that frame-shifted the protein coding sequence or removed the start codon. At some loci large insertions or repetitive sequences were introduced that could only be partially mapped by Sanger sequencing. To confirm these modifications, we also performed whole-genome sequencing using Oxford Nanopore MinION sequencing. This generated 1-3 reads across some of the modified loci that confirmed their genomic rearrangements but was not deep enough for precise base calling.

After the first round of mutagenesis, we generated a cell line (clone 25) that contained frame-shifting indel mutations in MPZL1, MPZL3, SCN3B, deletion of the start codon of SCN1B, and an in-frame deletion of MPZ (Figure 1C). Clone 25 was subject to a second round of mutagenesis, and this led to the generation of BeHAPe cells. BeHAPe cells contain additional frame-shifting indel mutations in SCN4B, JAML and EGFP, an in-frame deletion of SCN2B and a larger in-frame deletion of MPZ.

The impact of the gene disruptions at the protein level are shown in Figure 1D. The gene-edits completely disrupted the protein coding sequences of SCN1B, MPZL2, MPZL3, while the protein coding sequences of SCN3B, SCN4B and MPZL1 encode only a truncated N terminus. While the deletion in MPZ left the codons for the signal sequence (M1-A29) intact, it removed I30-E71, which encodes for a central β-strand in the Ig domain (Figure 1E). This deletion should render the Ig domain of MPZ misfolded and possibly subjected to ER associated degradation, as is the fate of MPZ mutants associated with Charot-Marie-Tooth disease (21). Similarly, SCN2B was deleted of V40-Y69, which encodes a central β-strand in the Ig domain and thus should also render the protein misfolded.

The edited genomic loci in BeHAPe cells are included in Supporting Information and are illustrated in Figure 2. In SCN1B, exon 1 had a 6 bp deletion that included the start codon. In SCN2B, exon 2 had an in-frame 90 bp deletion. In SCN3B, 1407 bp were deleted including a frame-shifting 95 bp from 3’ end of exon 2. In SCN4B, exon 1 had an in-frame 30 bp deletion and exon 2 had a frame-shifting 92 bp deletion. In, MPZ exon 2 had an in-frame 126 bp deletion. In MPZL1, a large 43,475 bp region that included portions of exon 1 and 2 was deleted and replaced with ∼ 427 bp. In MPZL2, a frame-shifting 32 bp from exon 1, which included the start codon, was replaced with 27 bp which kept the protein coding sequence out of frame. In MPZL3, ∼ 397 bp of repetitive sequences was inserted just after the encoded initiating Met. There was also a single bp insertion in exon 2, causing a frameshift mutation. In JAML, exon 2 had a frame-shifting ∼ 2888 bp insertion. The JAML locus could not be genotyped with Sanger sequencing but was genotyped using long read whole-genome sequencing. This revealed that the inserted DNA was a partial duplication of intron 2 and exon 3 followed by a reversed exon 3, intron 2 and exon 2.

**Figure 2.**
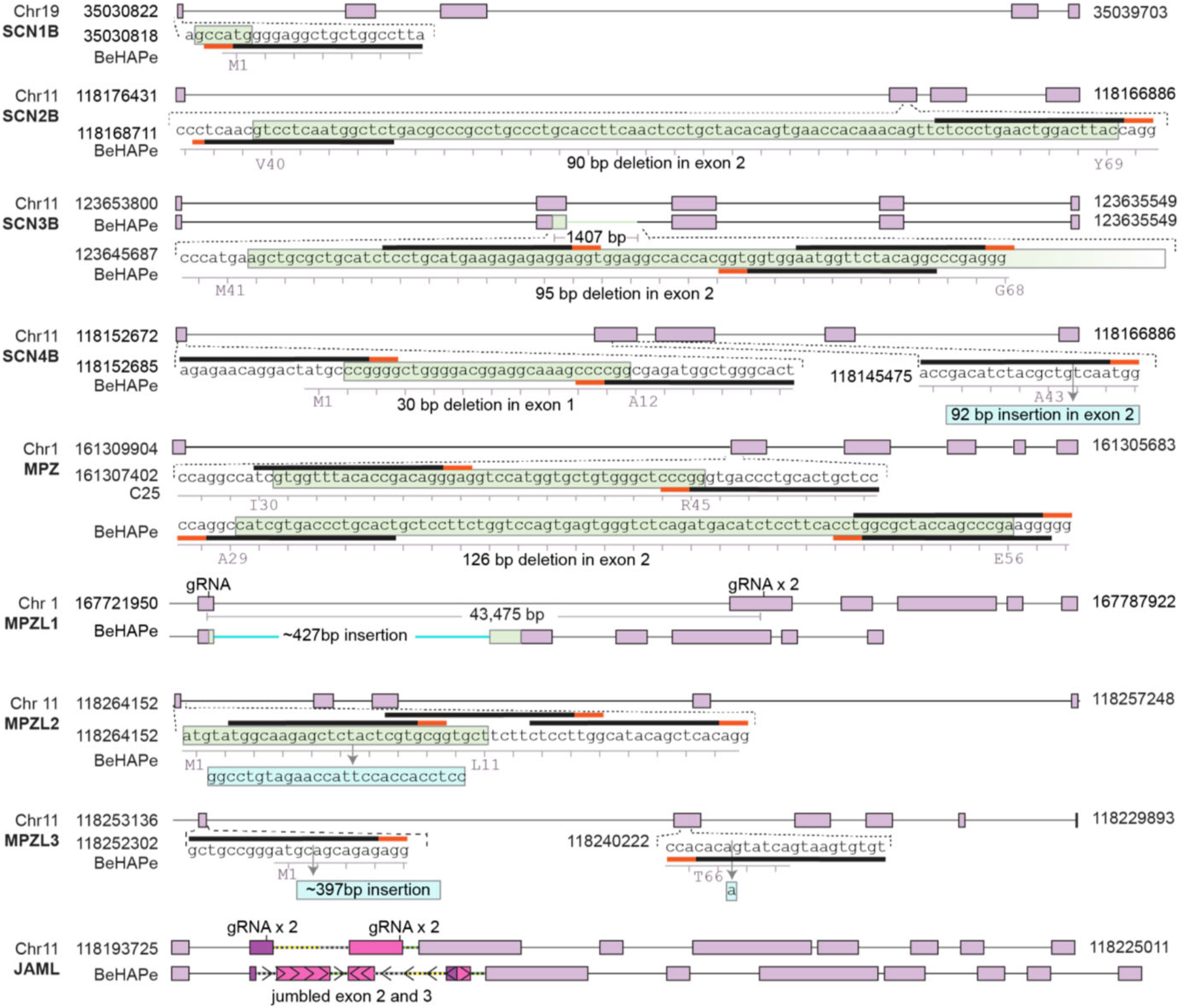
Mutations in β-subunits and their phylogenetic relatives in BeHAPe cells. Exons are drawn as purple boxes, deleted regions are colored green, and insertions are colored cyan. gRNA binding sites are indicated with a black and red bar, red bar indicates PAM sequence. The position of gene loci are derived from Genome Reference Consortium Human Build 38.

Surprisingly, we only recovered in-frame deletions of MPZ. In the first round of mutagenesis only a single clone (Clone 25) with an edited locus was recovered, and its locus had an in-frame deletion (Figure 2). In the second round of mutagenesis several clones with further edited MPZ were recovered, yet all had in-frame edits. This included three clones with in-frame deletions; one clone with a frameshift deletion, which was corrected with a second frameshift deletion that restored the wildtype reading frame; and two clones that had replaced sequences with new sequences that preserved the reading frame. All these deletions left the signal sequence intact, and the largest span of deleted residues was between I30-E71. These data suggest that this protein may be essential in haploid eHAP cells. If so, the essential function it serves must not require the extracellular Tg domain since this domain was disrupted in BeHAPe cells. The major known function of MPZ is in the compaction of myelin, a function irrelevant to eHAP viability. In addition, MPZ knockout mice are viable indicating MPZ is not essential in multiple cell types (23).

### Patch-clamping BeHAPe cells

We next sought to examine the electrophysiological properties of the cardiac Na_V_ channel Na_V_1.5 in BeHAPe cells, however, whole cell patch-clamp experiments were initially challenging. BeHAPe cells were small, exhibited a low transfection efficiency, and a plasma membrane that was prone to blebbing, which made it difficult to obtain tight giga-Ohm seals. This plasma membrane morphology was not observed in eHAP cells, which were easier to patch-clamp. These problems were partially mitigated through the following optimization steps.

First, while haploid BeHAPe cells (and parental eHAP cells) are small, diploid BeHAPe cells are larger and easier to patch-clamp (Figure 3). We obtained a diploid population by passaging haploid cells, which spontaneously diploidize, and sorting for diploid cells by FACS based on forward (FSC) and side-scatter (SSC) profiles (24). We confirmed that the magnitude of FSC and SSC corresponded to haploid and diploid eHAP cells by staining DNA content in live cells using Hoescht 33342 (Figure 3B). As Hoescht 33342 was toxic to cells, for routine sorting we set FSC and SSC gates using unstained reference populations.

**Figure 3.**
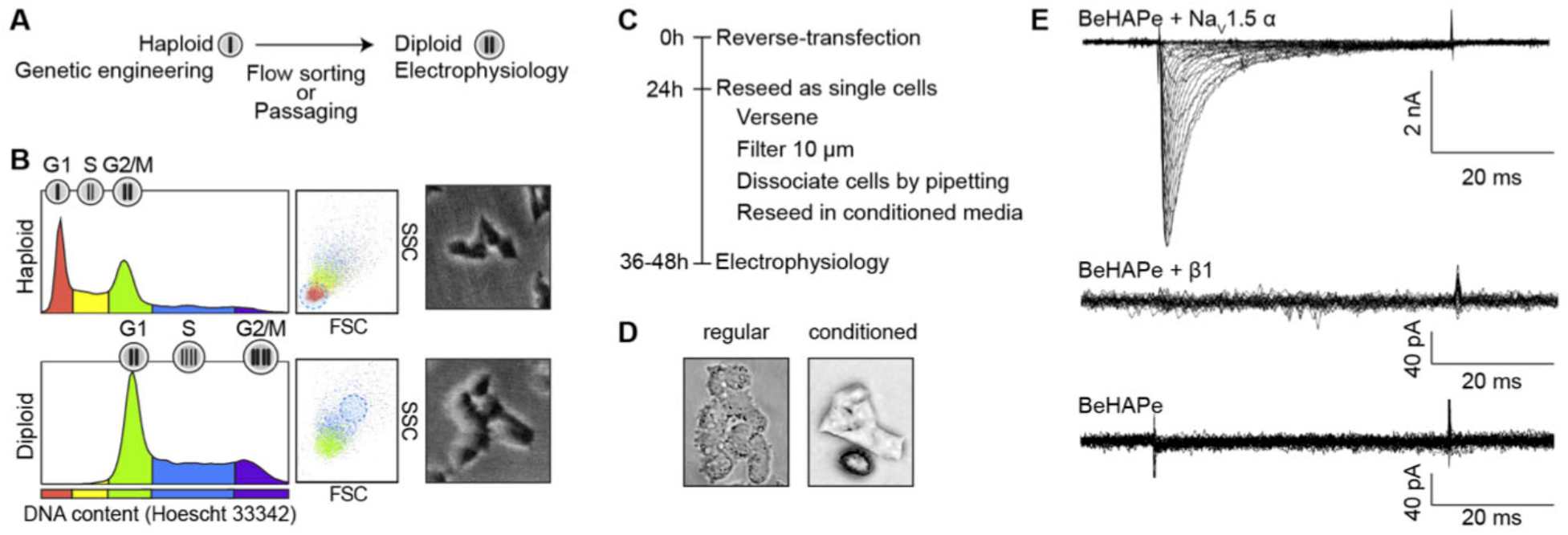
Preparation of BeHAPe cells for electrophysiology. **(A)** Haploid cells were used for gene-editing, while diploid cells were used for electrophysiology experiments. Haploid cells become spontaneously diploid after several passages. **(B)** Haploid or diploid BeHAPe cells can be enriched for by flow sorting based on cell size. BeHAPe cells were stained with Hoechst 33342 and DNA content was compared to forward (FSC) and side (SSC) scatter by flow cytometry. A micrograph of sorted haploid and diploid cells are shown. **(C)** Diploid BeHAPe cells were prepared for electrophysiology as follows. Cells were ‘reverse’-transfected and 24 h later reseeded as single cells. To obtain healthy single cells, cells were lifted with versene, resuspend in conditioned media, passed through a 10 μm mesh, and dissociated further with gentle pipetting. 36-48 h after transfection cells were phenotyped with electrophysiology. **(D)** BeHAPe cell membranes became blebby and incompatible with patch-clamping when resuspended in fresh media, however, remained smooth in conditioned media. **(E)** Sodium current recordings from BeHAPe cells un-transfected or transfected with β1 or Na_V_1.5 cDNA, as indicated.

Second, we achieved the highest expression of Na_V_1.5, with ∼ 30% transfection efficiency, in BeHAPe cells when they were ‘reverse’-transfected. Cells were reseeded as single cells 24 hours after transfection and patch-clamped 12-16 hours later, when there was peak expression of a tracer plasmid pmaxGFP (Figure 3C). Since the expression of transgenes was variable, we analyzed only cells with similar levels of the GFP tracer and current density. This selection approach however, precluded the use of peak current density as a proxy for trafficking of the a-subunit to the plasma membrane.

Finally, the plasma membrane of BeHAPe cells was prone to blebbing. The membrane blebbing was not suppressed by re-expression of β/MPZ subunits indicating a likely off-target effect. Plasma membrane blebbing could, however, be suppressed by growing BeHAPe cells in conditioned media and by avoiding sheer stress (Figure 3D). To reseed cells as single cells prior to electrophysiology, we reduced the amount of pipetting, and thus shear stress, by passing cells through a l0 μm filter, which is approximately the diameter of a cell, and seeding dissociated cells in conditioned media. During patch-clamp experiments, the cells retained good morphology in external solution for 30 minutes.

These methods allowed whole cell patch-clamp experiments on BeHAPe cells. Like eHAP cells, BeHAPe cells show no voltage dependent currents when transfected with an empty plasmid vector or β-subunit (Figure 3E). Robust currents are produced only after transfection with the α-subunit Na_V_1.5. Thus, this system can be used to reliably measure the electrophysiological properties of Na_V_ channels.

### Comparing electrophysiological effects of β1 on Na_V_1.5 in eHAP and BeHAPe cells

To determine whether eliminating the endogenous β/MPZ genes rendered the BeHAPe cells more sensitive to β-subunit modulation of Na_V_ channels, we compared the electrophysiological properties of Na_V_1.5 with and without β1 in eHAP cells and BeHAPe cells (Figure 4A and 4B). The eHAP cells revealed no β1-subunit effects on conductance-voltage curves (GV), steady-state inactivation curves (SST) or rates of inactivation compared to Na_V_1.5 expressed alone (Figure 4A). The data revealed that Na_V_1.5 without co-expressed β-subunits exhibited similar GV and SST in BeHAPe and eHAP cells, and the rate of inactivation was similar when analyzed at the V_1/2_ of the GV, but all were shifted in BeHAPe cells co-expressing β1 (Figure 4B). Thus, these results demonstrate that eliminating the endogenous β/MPZ proteins unmasks the effects of exogenous β-subunits on Na_V_1.5.

**Figure 4.**
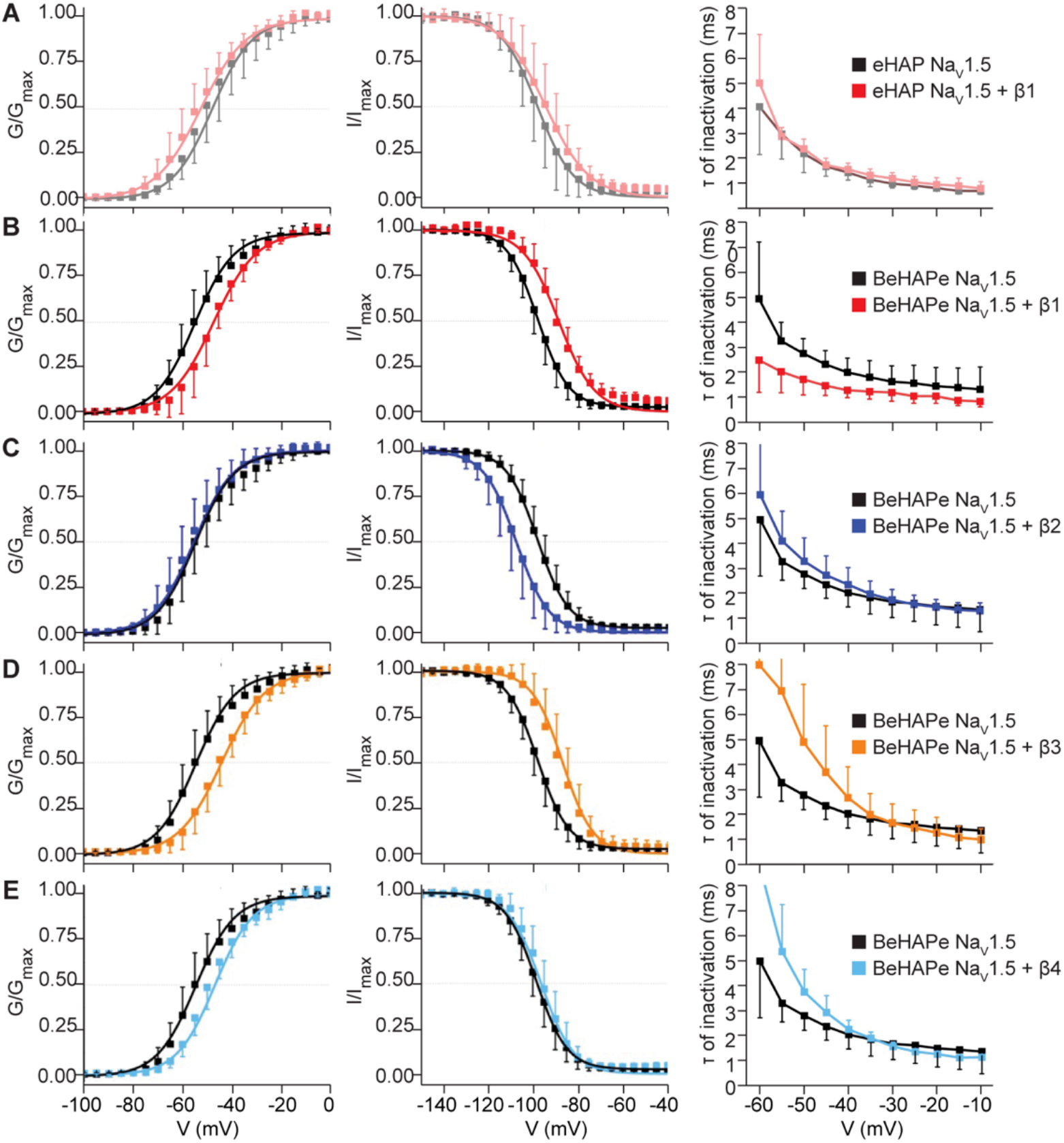
Biophysical properties of Na_V_1.5 when co-expressed with β-subunits part t. **(A)** Co-expression with β1 showed no effects on Na_V_1.5 biophysical properties in the parental eHAP cell line. Na_V_1.5 was expressed alone in eHAP cells and GV (n=8), SST (n=6) and the rate of inactivation (n=8) measured. When co-expressed with rat β1-V5-6xHTS, no significant change was found in GV V_1/2_ (n= 8, p=0.77), SST V_1/2_ (n=6, p=0.30), or the rate of inactivation at V_1/2_ of the GV (n=8, p=0. 4). **(B-E)** Co-expression with β1-4 modulates Na_V_1.5 biophysical properties in the BeHAPe cells. Na_V_1.5 was expressed alone in BeHAPe cells and GV (n=10), SST (n=9) and the rate of inactivation (n=10) measured. Plots for Na_V_1.5 alone in BeHAPe are identical across graphs. **(B)** Rat β1-V5-6xHTS co-expression shifted Na_V_1.5 GV V_1/2_ (6.7 3. mV, n=, p=0.0), SST V_1/2_ (9 mV, n=7, p=0.00), and rate of inactivation at V_1/2_ of the GV (−1.7 1.0 ms, n=, p=0.000 5). **(C)** Human β -HA co-expression shifted SST V_1/2_ (−9.9 3.1 mV, n=6, p=0.0019) but not GV V_1/2_ (n=6, p=0.33) or the rate of inactivation at V_1/2_ of the GV (n=6, p=0.30). **(D)** Human β3 co-expression shifted Na_V_1.5 GV V_1/2_ (.1 3.4 mV, n=7, p=0.003) and SST V_1/2_ (9.1 3.1 mV, n=6, p=0.0053), but not the rate of inactivation at V_1/2_ of the GV (n=6, p=0.34). **(E)** Human β4 co-expression shifted Na_V_1.5 GV V_1/2_ (5. 3.0 mV, n=6, p=0.045) but not SST V_1/2_ (n=6, p=0.46) or the rate of inactivation at V_1/2_ of the GV (n=6, p=0.30). **(A-E)** Data are plotted as the mean SD. For GV and SST plots, mean values were fitted to Boltzmann functions. For rates of inactivation plots, rates were derived from a single exponential function manually fit to individual traces.

### Electrophysiological effects of B/MPZ-proteins on Na_V_1.5 in BeHAPe cells

We next used BeHAPe cells to compare the individual effects of all β-subunits (B1-4) on the gating properties of the major cardiac α-subunit Na_V_1.5 (Figure 4 and Table 1). We also included MPZ in our analysis to gauge the capacity of β-subunit phylogenetic relatives to affect Na_V_1.5 gating (Figure 5 and Table 1). The voltage dependence of activation (GV), steady-state inactivation (SST), and rate of inactivation curves are plotted in Figure 4 and 5. A summary of the gating properties with individual replicates are displayed in Figure 5C and the values obtained are recorded in Table 1. Remarkably, the β-subunits and MPZ all modulated at least one electrophysiological property of Na_V_1.5.

**Table 1.** Gating properties of Na_V_1.5 in BeHAPe cells. Data are reported as mean ± SD. One-way ANOVA followed by Fischer’s Least Significance Difference was used to compare data to Na_V_1.5 alone in either eHAP (top) or BeHAPe (bottom); *p < 0.05, **p < 0.01. Data in this table were used to generate Figure 4 and Figure 5. Values of V_1/2_ for GV and SST as well as slopes were extracted from Boltzmann fits to mean values. Rates of inactivation were derived from a single exponential function manually fit to individual traces.

| Channel | GV | | | | SSI | | | | $\tau$ inactivation (at $V_{1/2}$) | | |
| --- | --- | --- | --- | --- | --- | --- | --- | --- | --- | --- | --- |
| | $V_{1/2}$ (mV) | $\Delta V_{1/2}$ (mV) | k (Slope) | N | $V_{1/2}$ (mV) | $\Delta V_{1/2}$ (mV) | k (Slope) | N | $\tau$ (ms) | $\Delta \tau$ (ms) | N |
| eHAP Nav1.5 | -49.8 ± 7.1 | -- | 8.2 ± 0.4 | 8 | -98.4 ± 7.3 | -- | 7.0 ± 0.4 | 6 | 3.0 ± 0.9 | -- | 8 |
| eHAP Nav1.5 + β1 | -53.4 ± 5.5 | 0.2 ± 3.3<br>($p = 0.77$) | 8.0 ± 0.3 | 8 | -94.6 ± 7.7 | 3.2 ± 3.4<br>($p = 0.30$) | 7.4 ± 0.3 | 8 | 2.9 ± 0.4 | -0.3 ± 1.0<br>($p = 0.839$) | 8 |
| BeHAPe Nav1.5 | -53.6 ± 5.6 | -- | 7.9 ± 0.4 | 10 | -97.8 ± 3.6 | -- | 6.8 ± 0.2 | 9 | 3.2 ± 0.6 | -- | 10 |
| BeHAPe Nav1.5 + β1 | -46.9 ± 4.9 | 6.7 ± 3.2*<br>($p = 0.020$) | 8.0 ± 0.4 | 8 | -89.0 ± 4.9 | 8.8 ± 2.9**<br>($p = 0.0088$) | 7.1 ± 0.3 | 7 | 1.5 ± 0.4 | -1.7 ± 1.0**<br>($p = 0.00025$) | 8 |
| BeHAPe Nav1.5 + β2 | -56.1 ± 6.1 | -2.5 ± 3.4<br>($p = 0.33$) | 8.2 ± 0.5 | 6 | -107.7 ± 6.5 | -9.9 ± 3.1**<br>($p = 0.0019$) | 6.4 ± 0.3 | 6 | 2.7 ± 0.7 | -0.5 ± 1.1<br>($p = 0.30$) | 6 |
| BeHAPe Nav1.5 + β3 | -44.8 ± 4.6 | 8.8 ± 3.1**<br>($p = 0.0030$) | 7.4 ± 0.9 | 7 | -88.7 ± 6.3 | 9.1 ± 3.1**<br>($p = 0.0053$) | 7.2 ± 0.5 | 6 | 3.7 ± 1.8 | 0.4 ± 1.5<br>($p = 0.34$) | 6 |
| BeHAPe Nav1.5 + β4 | -48.4 ± 3.4 | 5.2 ± 3.0*<br>($p = 0.045$) | 8.0 ± 0.3 | 6 | -95.4 ± 4.4 | 2.4 ± 2.8<br>($p = 0.46$) | 7.0 ± 0.4 | 6 | 3.7 ± 0.9 | 0.5 ± 1.2<br>($p = 0.30$) | 6 |
| BeHAPe Nav1.5 + β1 & β2 | -45.7 ± 4.3 | 7.9 ± 3.1**<br>($p = 0.0073$) | 8.1 ± 0.4 | 7 | -88.5 ± 5.4 | 9.1 ± 3.0**<br>($p = 0.0031$) | 7.2 ± 0.4 | 7 | 3.2 ± 1.1 | 0.0 ± 1.3<br>($p = 0.95$) | 7 |
| BeHAPe Nav1.5 + MPZ | -44.9 ± 6.5 | 8.7 ± 3.3**<br>($p = 0.0048$) | 7.4 ± 0.3 | 7 | -91.5 ± 4.7 | 6.3 ± 2.9*<br>($p = 0.035$) | 7.4 ± 0.3 | 6 | 3.0 ± 0.8 | 0.2 ± 1.2<br>($p = 0.65$) | 6 |
| eHAP Nav1.5 | -49.8 ± 7.1 | 3.8 ± 3.5<br>($p = 0.13$) | 8.2 ± 0.4 | 8 | -98.4 ± 7.3 | -0.6 ± 3.3<br>($p = 0.85$) | 7.0 ± 0.4 | 6 | 3.0 ± 0.9 | -0.2 ± 1.2<br>($p = 0.57$) | 8 |
| eHAP Nav1.5 + β1 | -53.4 ± 5.5 | 0.2 ± 3.3<br>($p = 0.80$) | 8.0 ± 0.3 | 8 | -94.6 ± 7.7 | 3.2 ± 3.4<br>($p = 0.44$) | 7.4 ± 0.3 | 8 | 2.9 ± 0.4 | -0.3 ± 1.0<br>($p = 0.21$) | 8 |

**Figure 5.**
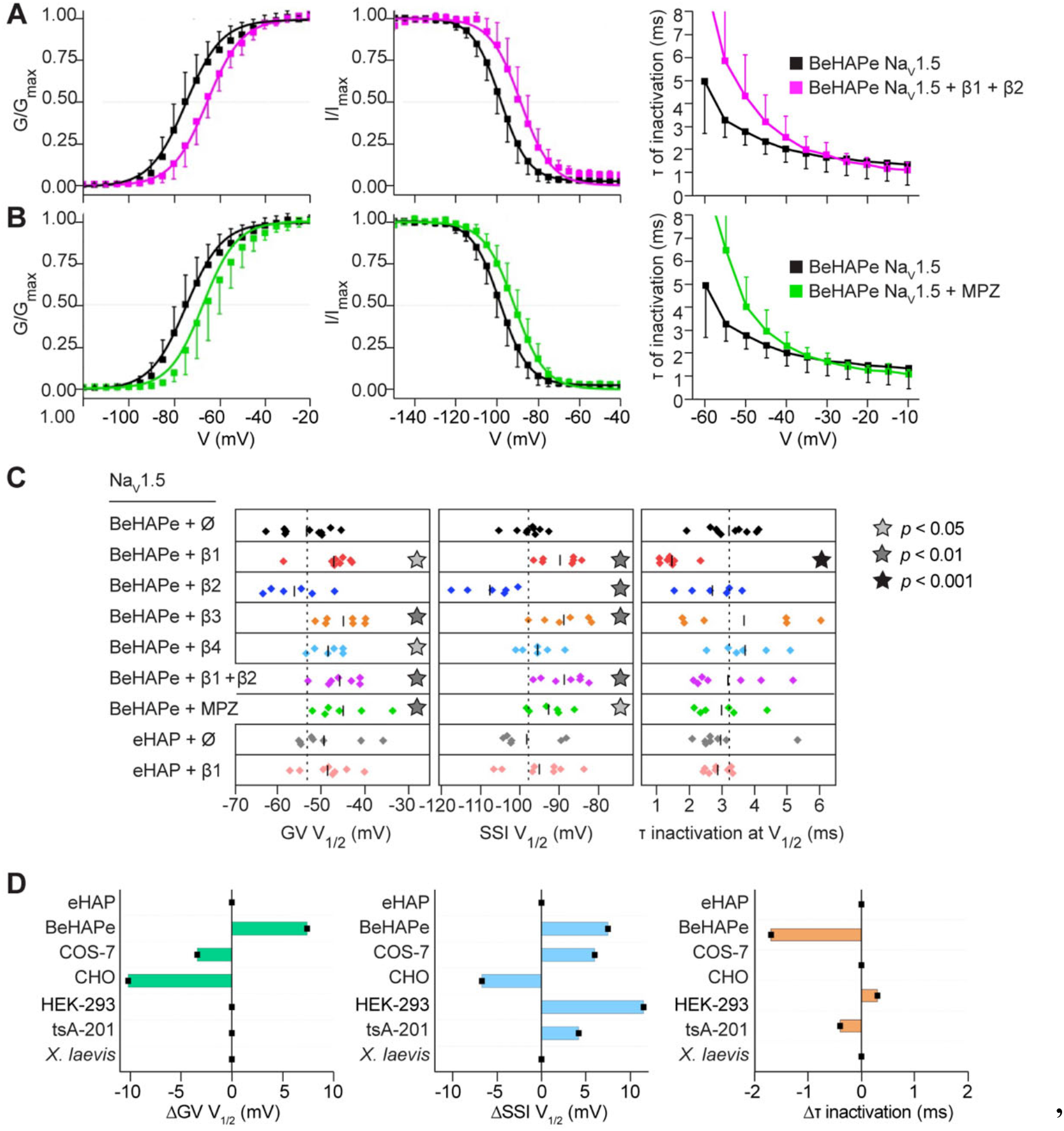
Biophysical properties of Na_V_1.5 when co-expressed with β-subunits part 2. **(A)**Simultaneous co-expression of both β1 and β2 shifted Na_V_1.5 GV V_1/2_ (+7.9 ± 3.1 mV, n=7, p=0.0073) and SST V_1/2_ (+9.1 ± 3.0 mV, n=7, p=0.0031) but not the rate of inactivation at V_1/2_ of the GV (n=7, p=0.95). **(B)**Human MPZ co-expression shifted Na_V_1.5 GV V_1/2_ (+8.7 ± 3.3 mV, n=6, p=0.0048), SST V_1/2_ (+6.3 ± 2.9 mV, n=6, p=0.035), but not the rate of inactivation at V_1/2_ of the GV (n=6, p=0.65). **(A**,**B)** Data for Na_V_1.5 alone in BeHAPe cells (n=10 for GV, n=9 for SST, n=8 for rate of inactivation) are identical to plots in Fig. 4. Data are plotted as the mean ± SD. For GV and SST plots, mean values were fitted to Boltzmann functions. For rates of inactivation plots, rates were derived from a single exponential function manually fit to individual traces. **(C)** Summary of the gating properties of Na_V_1.5 alone or co-expressed with the indicated β/MPZ subunits. Shown are GV V_1/2_, SST V_1/2_, τ of inactivation at V_1/2_ of the GV derived from the same experiments in Fig 4 and Fig 5 A and B. Data points are individual replicates. Black bars represent mean values, which are reported in Table 1. Star indicates values were significantly different compared to Na_V_1.5 alone in BeHAPe cells as determined by a one-way ANOVA followed by Fischer’s LSD. **(D)** Comparison of the magnitude of β1 modulation of Na_V_1.5 recorded in eHAP and BeHAPe cells to previously reported values in other heterologous systems. Electrophysiological properties are plotted as the difference between Na_V_1.5 alone and Na_V_1.5 co-expressed with β1. Values are reported in Tables 1 & 2

Co-expression with β1 in BeHAPe cells shifted the midpoint of the GV and SST curves of Na_V_1.5 to more positive potentials (+6.7 ± 3.2 mV for GV, +8.8 ± 2.9 mV for SST) without altering the slope of these curves (Figure 4B). β1 also increased the exponential rate of fast inactivation of Na_V_1.5 and was the only β-subunit to affect this property (−1.7 ± 1.0 ms) (Figures 4B, 5D).

β2 was the only β-subunit to shift the midpoint of the SST curve to more negative potentials (−9.9 ± 3.1 mV), and it did not shift the GV curve or the slope of either curve (Figure 4C and Table 1). The effect of B3 on GV and SST were almost identical to β1, with β3 shifting the midpoint of the GV and SST curves to more positive potentials (+8.1 ± 3.4 mV for GV and +9.1 ± 3.1 mV for SST) without altering slope or the rate of fast inactivation (Figure 4D and Table 1). Unlike the other β-subunits, β4 had no effect on the SST curve but did shift the midpoint of the GV curve to more positive potentials (+5.2 ± 3.0 mV shift in GV) (Figure 4E).

Since β1 and β2, when individually expressed, had opposing effects on Na_V_1.5 gating, we examined the effect on Na_V_1.5 gating properties when both were expressed simultaneously (Figure 5A). Tn contrast to the effects of β1, expressing both β1 and β2 did not significantly change inactivation rates, but did shift the midpoint of the GV and SST curves to more positive potentials (+7.9 ± 3.1 mV shift in GV, +9.1 ± 3.0 mV shift in SST) (Figure 5A and Table 1). These data indicated the effects of β1 override those of β2 on equilibrium gating, but the presence of β2 nullified the effects of β1 on inactivation kinetics.

We were surprised to find substantial effects on Na_V_1.5 gating by MPZ, which, like β1 and β3, shifted the midpoint of the GV and the SST curves to more positive potentials (+8.7 ± 3.3 mV for GV and +6.3 ± 2.9 mV for SST) but, as with β2-4, did not affect rates of fast inactivation (Figure 5B). Overall, these data demonstrate that each of the β-subunits and their phylogenetic relative MPZ have unique effects on Na_V_1.5 gating parameters, and that BeHAPe cells can be used as platform for revealing these effects.

## Discussion

Na_V_s produce electrical signals in excitable cells, and dysfunction in the channel’s pore-forming α-subunit or auxiliary β-subunits, are associated with a range of conduction diseases (7,25-28). β-subunits modulate the gating properties of α-subunits (2). Despite its importance, this function is difficult to study with currently available expression systems. Notably, the effects of β-subunits vary across expression systems as exemplified in Table 2 and Figure 5D (8,9,11,29-40). We hypothesized that the effects of ectopically expressed β-subunits could be better resolved in an expression system without endogenous β-subunits and their phylogenetic relatives, the MPZ-related proteins. Tn support of this hypothesis, we found that eliminating β/MPZ genes from eHAP cells sensitized Na_V_1.5 to modulation by ectopically-expressed B1. We used the newly generated BeHAPe cells, to evaluate the effects of β1-4 on the electrophysiological properties of the main cardiac volage-gated sodium channel, Na_V_1.5. We observed that each of the β-subunits uniquely modulated the Na_V_1.5 channel and the effects we obtained were generally different than observed in previous studies. These findings demonstrate the benefit of using expression systems devoid of endogenous β-subunits in basic studies on α/β interactions and may also be beneficial in studies on Na_V_ pharmacology and disease-causing mutations.

**Table 2.** Gating properties of Na_V_1.5 in previous studies. Values are presented as mean ± SEM. Data in this table were used to generate Figure 5D.

| Expression System | Channel | GV | | SSI | | $\tau$ inactivation (ms at -40mV) | | Publication |
| --- | --- | --- | --- | --- | --- | --- | --- | --- |
| | | $V_{1/2}$ (mV) | $\Delta V_{1/2}$ (mV) | $V_{1/2}$ (mV) | $\Delta V_{1/2}$ (mV) | $\tau$ (ms) | $\Delta\tau$ (ms) | |
| *X. laevis* | Nav1.5 | -40.6 ± 1.2 | | -69.2 ± 0.8 | | 0.7 ms | | (43) |
| | Nav1.5 + β1 | -40.6 ± 1.9 | NS | -70.5 ± 0.8 | NS | 0.7 ms | NS | |
| *X. laevis* | Nav1.5 | -18.6 ± 4.2 | | -65.3 ± 0.9 | | NR | | (53) |
| | Nav1.5 + β1 | -23.3 ± 2.0 | NS | -65.9 ± 0.7 | NS | NR | | |
| | Nav1.5 + β3 | -21.3 ± 2.4 | NS | -60.7 ± 1.3 | 5.2 | NR | | |
| tsA-201 | Nav1.5 | -25.6 ± 1 | | -77.1 ± 0.5 | | NR | | (9) |
| | Nav1.5 + β1 | -24.9 ± 1 | NS | -72.9 ± 1.0 | 4.2 | NR | | |
| | Nav1.5 + β2 | -23.2 ± 1 | 2.4 | -77.1 ± 1.1 | NS | NR | | |
| | Nav1.5 + β1 + β2 | -24.3 ± 2 | NS | -74.0 ± 1.0 | 3.1 | NR | | |
| tsA-201 | Nav1.5 | -35.4 ± 1.2 | | -88.2 ± 0.9 | | NR | | (44) |
| | Nav1.5 + β1 | -34.7 ± 1.2 | NS | -81.8 ± 0.9 | NS | NR | | |
| HEK-293T | Nav1.5 | -37.8 ± 1.6 | | -95.2 ± 2.9 | | 2.3 | | (45) |
| | Nav1.5 + β1 | -36.7 ± 0.6 | NS | -90.9 ± 2.1 | 4.3 | 1.9 | -0.4 | |
| HEK-293T | Nav1.5 | -43.78 ± 1.28 | | -78.77 ± 1.11 | | 0.68 (-20mV) | | (11) |
| | Nav1.5 + β4 | -43.69 ± 1.61 | NS | -82.28 ± 0.74 | -3.5 | 0.77 (-20mV) | NS | |
| HEK-293 | Nav1.5 | NR | | -70.2 ± 1.3 | | NR | | (8) |
| | Nav1.5 + β1 | -22 | | -58.7 ± 1.2 | 11.5 | 1.1 ms | | |
| CHO | Nav1.5 | -38.7 ± 0.5 | | -86.9 ± 1.5 | | NR | | (42) |
| | Nav1.5 + β1 | -48.9 ± 0.8 | -10.2 | -93.6 ± 0.8 | -6.7 | NR | | |
| | Nav1.5 + β2 | -38.0 ± 0.6 | NS | -80.2 ± 0.6 | 6.7 | NR | | |
| CHO-K1 | Nav1.5 | -16.2 ± 1.0 | | -61.7 ± 1.9 | | 2.0 ms | | (46) |
| | Nav1.5 + β1 | -22.2 ± 1.8 | -6 | -68.7 ± 2.5 | -6.7 | 2.3 ms | 0.3 | |
| COS-7 | Nav1.5 | -33.5 ± 0.3 | | -95.8 ± 0.3 | | 1.2 ms | | (40) |
| | Nav1.5 + β1 | -36.1 ± 0.6 | -3.4 | -89.8 ± 0.2 | 6 | 1.1 ms | NS | |

The study of Nav channels in heterologous systems remains a valuable approach for the precise measurement of channel electrophysiological properties. By removing the channel from its native context, currents generated only by a channel-of-interest can be studied. However, the ability to study β-subunit effects on Na_V_ channels may be limited in currently available systems since they express endogenous β-subunits and their phylogenetic relatives. When comparing the results reported here with similarly conducted studies in HEK-293, COS-7, and CHO cells, and *X. laevis* oocytes, we found the β-subunit effects to be of different magnitude and, in some cases, different relative directions in BeHAPe cells (Figure 5D and Table 2) (8,9,11,29-40). Notably the effects of β1 on the voltage-dependence of activation observed in BeHAPe cells have not been observed in *X. laevis* oocytes or HEK-293 cells, and the effects on the voltage-dependence of inactivation have not been observed in *X. laevis* oocytes (29,31,32,41).Furthermore, the effect of β1 on the rate of Na_V_1.5 inactivation was large compared to other systems. Na_V_1.5 produced a voltage-gated sodium current with a rate of fast inactivation that was relatively slow compared to previously published values in other expression systems (2.0 ms at -40mV). However, when co-expressed with β1, the inactivation rate increased to values similar to those seen in other systems (1.3 ms at -40mV). These results suggest that the BeHAPe cells provide a sensitive platform for revealing B-subunit effects on electrophysiological properties of Na_V_ channels. Our findings show the complications that endogenous expression of β-subunits can pose. Notably all the β-subunits regulated Na_V_1.5 channel, as did a structurally related protein like MPZ. Thus, to avoid interference from endogenous β-subunits, the entire family should be removed.

There are many Na_V_ channels for which the precise electrophysiological properties are sought after, and thus would be best resolved in a clean β/MPZ null background. These include the nine other α-subunits, which may have different responses to the β-subunits, and clinical variants of α and β-subunits. However, phenotyping numerous α/β channels in BeHAPe cells is difficult because the cells are not compatible with high-throughput screening. BeHAPe cells are not as robust as standard cell lines and require careful preparation for transfection and to avoid membrane blebbing. For these studies it would be beneficial to engineer a more robust cell line, which might be achieved by deleting B/MPZ proteins from, for example, HEK-293 cells. This heterologous expression system could help characterize the electrophysiological properties of Na_V_ channels and complement studies in differentiated pluripotent stem cells or primary cells that study Na_V_ channels in their more complex native environment.

Despite the clear association of Na_V_1.5 with β-subunits, the structural basis for association and modulation of the Na_V_1.5 α-subunit remains unknown (42,43). We found that the β-subunits each elicited a unique combination of effects on the GV, SST, and kinetics of inactivation on Na_V_1.5. Tnterestingly β-subunits most similar in sequence exerted similar effects. β1 and β3 both shifted the GV and SST to more positive voltages, whereas β2 and β4 did not. MPZ, which is similar in sequence to β1 and β3, also shifted the GV and SST to more positive potentials. Although not a physiological interactor of Na_V_1.5, MPZ can be used as a structural tool for gaining insight into the molecular basis of the differential effects of the β-subunits. We also found that the combination of β1 and β2 modulated Na_V_1.5 in ways that were distinct from the effects of individual subunits (Table 1). Some Na_V_ α-subunit isoforms are known to be capable of binding multiple β-subunits simultaneously (44,45). Na_V_1.5 may have this property as well, allowing it to form a hybrid complex with different properties. Further experiments in a cell background such as the BeHAPe cells now enable the nuanced regulatory effects of β-subunits to be dissected.

The unique effects of β1-β4 on Na_V_1.5 gating properties could play a major role in cardiac rhythm. Although the specific function of β-subunits in heart are not defined, they are physiologically important as mutations in the subunits are associated with arrhythmias, and knock out of β-subunits (β1-β3) in mice cause cardiac arrythmia (12-14). Tntriguingly the β-subunits are reported to occupy different subcellular regions in cardiac myocytes and concentrate in different regions of the heart (9,46-49). It seems likely that β-subunits operate *in vivo* to fine-tune heart physiology by modulating Na_V_1.5 gating at different locales. Another intriguing property of Na_V_1.5 revealed here is that the inactivation kinetics of the α-subunit are accelerated by only β1. This property may be linked to forms of cardiac dysfunction that are accompanied by a persistent sodium current caused by altered inactivation kinetics (50-52). If so, altered association with β-subunits within a subpopulation of Na_V_1.5 complexes might contribute to such a late current. We also observed that MPZ regulated Na_V_1.5, and although this is unlikely to be a physiological interaction, given that MPZ is not expressed in the heart (53), there is a possibility that MPZ family proteins do regulate Na_V_ channels physiologically. For example, MPZL2 and MPZL3 are expressed in cardiac myocytes and SCN5A is expressed in other cells besides cardiac myocytes (proteinatlas.org)(54). The broader expression pattern of the B/MPZ-subunit family, and thus its potential for altering the behavior of other Na_V_ isoforms, remains to be explored.

Our strategy for studying the β-subunit family can be employed for characterizations of other gene families. Genes within a family, such as Na_V_ β/MPZ-subunit family, can often compensate for each other due to their structural similarity. In these circumstances, a gene-of-interest might only reveal its full range of activity in an expression system devoid of all compensating isoforms. While deletion of multiple genes from an expression system was previously time consuming, we show that it can be done relatively efficiently in haploid eHAP cells. The disadvantage of transfecting multiple gRNAs simultaneously is that recovered clones may have off-target mutations, such that BeHAPe cells may not be precisely congenic with its parent. For this reason, a gene-of-interest should always be characterized by reintroducing it into the new cell line, rather than by comparing the mutant cell line to the wild-type parental cells.

Besides Na_V_1.5, there are nine other α-subunits, and each may be modulated differently by the β-subunits. There are also disease-causing mutations in α and β-subunits that may affect channel properties. Our studies in BeHAPe cells demonstrate that the consequences of these interactions are best defined in a clean β/MPZ null background. However, as the BeHAPe cells are only suited to low throughput analysis, a systematic high-throughput electrophysiology analysis would greatly benefit from the engineering of a robust B/MPZ null cell line.

### Experimental Procedures

#### Molecular Biology Reagents

Plasmids and oligonucleotides used in this study are described in Supporting Information (Supplemental Table 1). All enzymes targeting DNA and RNA were obtained from New England Biolabs (Ipswitch, MA).

### Cell culture

Human haploid eHAP cells (Cat # C669, Horizon Discovery, Cambridge MA) were cultured in Iscove’s Modified Dulbecco’s Medium (IMDM: Gibco, Billings, MT) supplemented with 10% FCS. Cells were passaged every 48-72 h. As noted previously, eHAP cells can diploidize and make aneuploid populations (24). Thus, eHAP cells and their derivates were enriched for haploid cells using flow-actuated cell sorting (FACS) using size (FSC: forward scatter; SSC: side scatter) as the sorting parameter. Haploid-enriched populations of eHAP cells were used for gene engineering, whereas diploid cells were used for electrophysiology experiments. For transfection, Lipofectamine 3000 (Invitrogen/Thermofisher, Waltham, MA) was used in a reverse-transfection scheme as previously described (55) using manufacturer’s instructions for making DNA-Lipofectamine particles. Some culture conditions required conditioned media, which was made as follows: BeHAPe cells were seeded onto a 100 mm dish, allowed to become confluent, after which they were grown for an additional week. The media was then harvested and passed through a 0.22 μm filter prior to storage at 4°C until used.

### Generation of BeHAPe cells

A Flp Recombination Target site (FRT) was first integrated into eHAP cells. eHAP cells were transduced with lentivirus carrying the pQCXTP FRT GFP-Neo^R^ (pPL6490) vector. pPL6490 encodes an FRT site, GFP and neomycin-resistance. pPL6490 was derived from pQCXTP (56), a bicistronic retroviral expression vector that originally conferred puromycin-resistance. pPL6490 was packaged into lentivirus in HEK293 cells using the previously described lentivirus packaging system (57). Transduced eHAP cells were selected for by isolating neomycin-resistant colonies that express GFP. From these clones we identified eHAP-FRT. The FRT integration site in eHAP-FRT was defined by isolating the integrated plasmid DNA and the genomic DNA flanking the integration site. This was achieved by first digesting genomic DNA with the restriction enzymes HindTTT-HF, EcoRV-HF, HpaT, ApaT, BamHT-HF, XhoT, StuT, as well as RNAaseA and Klenow fragment in the presence of 1 mM dNTPs. Digested genomic DNA was ligated with T4 DNA ligase and electroporated into SURE bacterial cells (Agilent, Santa Clara, CA).

### Plasmids from ampicillin-resistant colonies were Sanger sequenced to define the FRT insertion described

To disrupt genes encoding the β/MPZ family, two rounds of CRTSPR-Cas9 gene-editing were performed on eHAP-FRT cells. Multiple gRNA sequences that targeted the N-terminal signal sequences or the exofacial Tg-like domains were used per gene. Tn Round 1, eHAP-FRT cells were transiently transfected with pU6-(BbsT)_CBh-Cas9-T2A-mCherry (58) encoding *S. pyogenes* Cas9 and mCherry, and nine plasmids derived from pCLTP dual SFFV ZsGreen (Transomics, Huntville AL) that encoded gRNAs against MPZ, MPZL1, MPZL3, SCN1B, SCN3B (Supplemental Table 1). After 48 h, ZsGreen and mCherry double-positive cells were isolated by FACS as described below, and serially diluted into 100 mm plates and allowed to form colonies. Colonies were isolated using trypsin-soaked discs as previously described (30) and assessed for ploidy using propidium iodide staining. Haploid populations were genotyped as described below. One clone (clone 25) had frame-shifting indel mutations in all targeted genes, although MPZ had an in-frame deletion. Clone 25 cells were FACS-sorted for haploid cells and then subjected to another round of CRTSPRICas9 mediated mutagenesis targeting SCN2B, SCN4B, JAML, MPZ and EGFP at the FRT locus. Here, gRNA encoding sequences were cloned into the BbsT site downstream of the U6 promoter of pU6-(BbsT)_CBh-Cas9-T2A-mCherry to generate 16 different plasmids. Clone 25 cells were transiently transfected with these 16 plasmids, and 48 h after transfection, cells positive for mCherry were isolated by FACS, serially diluted, and allowed to form colonies after plating. Clones were isolated on cloning discs, and after expansion were assessed for ploidy using propidium iodide staining and genotyped. One clone, BeHAPe cells, had frame-shifting indel mutations in all targeted genes, except for MPZ and SCN2B, which acquired a large in-frame deletion. BeHAPe cells were FACS-sorted for haploid cells before storage.

### Genotyping of eHAP-derived cells

Genomic DNA was harvested using proteinase K and phenol-chloroform extraction (59). The CRTSPR-targeted genetic loci were PCR amplified from genomic DNA using NEBNext High-Fidelity 2X PCR Master Mix and the primer pairs in Supplemental Table 1. PCR products were purified, and Sanger sequenced using the primers also listed in Supplemental Table 1. We also performed whole genome sequencing using an Oxford Nanopore MinION equipped with a version 10.41 flow cell. Reads were mapped using miniMAP2 using the default parameter for Oxford Nanopore data (60). Sanger sequences of PCR amplicons and Oxford Nanopore DNA sequences of edited loci are included in Supplementary Sequence Data as are the annotated sequences of the edited loci.

### Ploidy analysis and FACS

The ploidy of eHAP cells was determined by propidium iodide staining and analyzed on a Becton Dickinson LSR II flow cytometer. Cells were trypsinized, washed twice with PBS, lysed and stained using Nicoletti buffer (0.1% sodium citrate, 0.1% Triton X-100, 0.5 unit/mL RNase A, 20 units/mL RNase T1, 50 μg/mL propidium iodide). Haploid eHAP cells were used for reference. Ploidy was also assessed by Hoescht 33342 staining. Hoescht 33342 (5 ug/mL) was applied to live eHAP cells for 10 min, cells were then trypsinised and fluorescence intensity measured on a Becton Dickinson Aria II. Cells showed strong correlation between cell size and Hoescht intensity allowing the former parameter to be used to enrich haploid cells via cell sorting as previously described (24).

For cell sorting experiments, a Becton Dickinson Aria II equipped with a 130 μM diameter nozzle was used. Haploid or diploid cells were enriched by gating based on FSC and SSC. Reference populations were used to define the gates. Cells transiently expressing mCherry and/or ZsGreen were sorted for by using gates that captured the brightest 0.3 % of cells.

### Preparation of cells for electrophysiology experiments

Electrophysiology experiments were performed on diploid BeHAPe cells using ‘reverse’ transient transfection of the plasmids listed in Supplemental Table 1. All constructs were grown in DH5α cells (NEB #2987H), prepared using PureLink HiPrep plasmid preparation kits (Invitrogen/Thermofisher, Waltham, MA), followed by sequencing of the reading frame and promoter. All constructs expressed their respective open-reading frame via the CMV promoter. Cells were transfected with plasmids expressing Na_V_1.5-V5, pmaxGFP expressing *Pontellina plumata* GFP (Amaxa Biosciences, Cologne, Germany), and either empty vector (pcDNA3.1) or plasmid expressing a B/MPZ family member using a ratio of 2:1:2, respectively. 24 h after transfection, cells were dissociated from tissue culture dish using Versene (Gibco, Billings, MT), washed and resuspended in conditioned IMDM with 10% FCS. Cells were then immediately passed through a 10 μM nylon net filter (Milipore) and dispersed further by gentle pippetting before plating in 35 mm corning cell culture dishes (SKU: CLS430165, Corning, Tewksbury, MA). 12-24h after reseeding cells, electrophysiology experiments were performed.

### Whole-cell voltage patch clamp

Ionic currents through Na_V_1.5 (SCN5A) channels expressed in BeHAPe cells were recorded using whole-cell patch on Axon Axopatch 200B amplifiers (Molecular Devices, San Jose, CA). Data were collected and analyzed with pClamp11/Clampfit11 (Molecular Devices, San Jose, CA) and Origin software (OriginLab, Northhampton, MA). Glass microelectrodes had resistances of 1.5 − 2 M n. Internal solution consisted of 105 mM CsF, 33 mM NaCl, 10 mM HEPES, 10 mM EDTA, pH-adjusted to 7.3 with CsOH. External solution contained 150 mM NaCl, 2 mM KCl, 1.5 mM CaCl_2_, 1 mM MgCl_2_, 10 mM HEPES, pH-adjusted to 7.4 with NaOH. Included recordings had currents between 1 nA and 12 nA, access resistance of <6 MΩ, and compensated series resistance of >90%. All cells were recorded 36-48 h post-transfection. Data from at least 3 separate transfections constituted each dataset. Cells of similar size were analyzed by recording only from cells with whole-cell capacitance between 0.5 − 1.0 pF. Data were sampled at 20 kHz and filtered at 5 kHz. Leak currents were subtracted with the p/8 protocol (61), except in the case of the β-subunit only control conditions wherein the minute leak current was not subtracted to ensure integrity of control measurements. The Conductance-Voltage relationship (GV) curves were determined by dividing the measured current amplitude at a given test voltage by the driving force. GV and SSI measurements were averaged, and the means fit to a Boltzmann function confined between 0-1 to produce experimental values. For rates of inactivation, all data were P/8-subtracted and currents were fit to a single exponential with the function y=A+A_0_exp^−t/t^, with the bounds of the fit currents set manually to include the fast inactivating portion of the trace. Comparison of these rates was performed at the V_l/2_ of the GV, respective to experimental condition, to ensure equivalence of analysis regardless of shifts in GV relationships. Multiple authors conducted blinded exponential fits ensure fit integrity. Example trace data for different conditions are provided in Supporting Information (Figure S1)

### Statistical Analysis

Statistical analyses were carried out using Origin Lab. Statistical significance was assessed with a one-way ANOVA with Fisher’s Least Significance Difference (LSD) *post-hoc* test without correction for multiple comparisons. Data are presented as mean ± SD. Individual values are reported in Supplemental Table 2 in Supporting Information.

## Supporting information

Supplementary Sequence Data

## Data Availability

BeHAPe cells are available upon request from Robert Piper or Chris Ahem, University of Iowa. Annotated sequence maps of BeHAPe genes that were mutated and the key sequence data used to determine the BeHAPe genotype are included in Supporting Information. All data used to generate the electrophysiology graphs and tables are also provided in Supporting Information.

## Supporting Information

This article contains supporting information.

## Acknowledgements

This work was supported by NIH-R01 GM106568 to CAA, and NIH RO1GM058202 to RCP. AYM was supported by ADA postdoctoral fellowship. CAA and RCP were supported by the Roy J. Carver Charitable Trust.

## Author Contributions

AYM: Conceptualization, Methodology, Investigation, Visualization, Writing-Original draft preparation, Writing-Reviewing and Editing. CJC: Conceptualization, Investigation, Visualization, Writing-Original draft preparation. CAA: Supervision, Conceptualization, Writing-Reviewing and Editing, Funding acquisition. RCP: Supervision, Conceptualization, Writing-Reviewing and Editing, Funding acquisition.

---

**Supplemental Figure 1.**
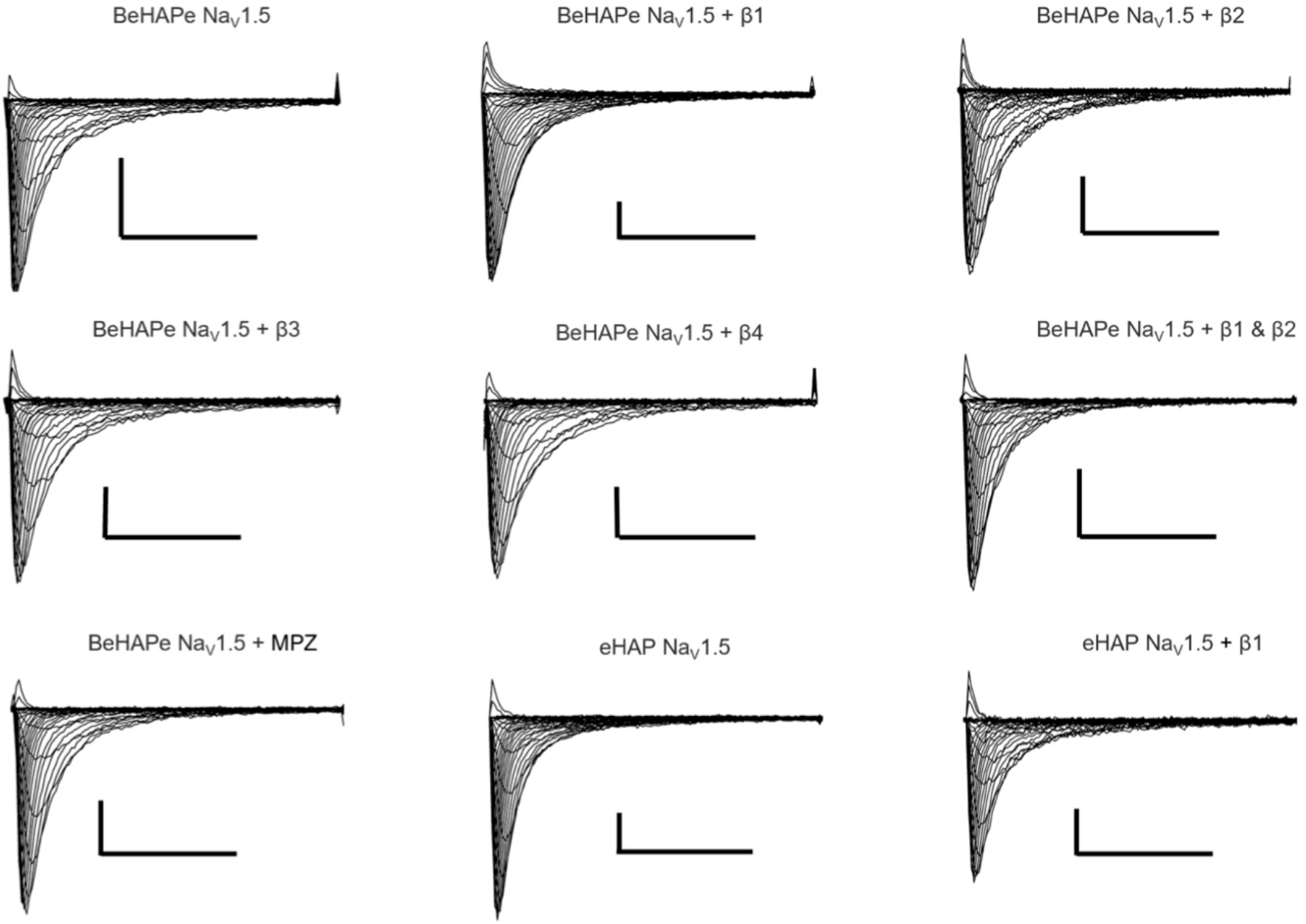
Representative trace families of each experimental condition. All scale bars are 1nA and 10ms.

